# Discovery of aphid-transmitted Rice tiller inhibition virus from native plants through metagenomic sequencing

**DOI:** 10.1101/2022.11.04.515094

**Authors:** Wenkai Yan, Yu Zhu, Wencheng Liu, Chengwu Zou, Bei Jia, Zhong-Qi Chen, Yanhong Han, Jianguo Wu, Dong-lei Yang, Baoshan Chen, Rongbai Li, Shou-Wei Ding, Qingfa Wu, Zhongxin Guo

## Abstract

A major threat to rice production is the disease epidemics caused by insect-borne viruses that emerge and re-emerge with undefined origins. It is well known that some human viruses have zoonotic origins from wild animals. However, it remains unknown whether native plants host new endemic viruses with spillover potential to rice (*Oryza sativa*) as emerging pathogens. Here, we discovered rice tiller inhibition virus (RTIV), a novel RNA virus species, from colonies of Asian wild rice (*O. rufipogon*) in a genetic reserve by metagenomic sequencing. We identified the specific aphid vector to transmit RTIV and found RTIV would cause low-tillering disease in rice cultivar after transmission. We further demonstrated that an infectious molecular clone of RTIV initiated systemic infection and causes low-tillering disease in an elite rice variety after Agrobacterium-mediated inoculation or stable plant transformation, and RTIV can also be transmitted from transgenic rice plant through its aphid vector to cause disease. Finally, global transcriptome analysis indicated that RTIV may disturb defense and tillering pathway to cause low tillering disease in rice cultivar. Thus, our results show that new rice viral pathogens can emerge from native habitats, and RTIV, a rare aphid-transmitted rice viral pathogen from native wild rice, can threaten the production of rice cultivar after spillover.

## Introduction

Rice (*Oryza sativa*) is one of the most important staple crops in the world [1]. Much is known about the mechanisms of key breeding traits of rice such as yield and disease resistance [2–5]. However, most of the current conventional and hybrid rice varieties remain vulnerable to the diseases caused by diverse families of viruses [6]. All but one of the viruses known to infect rice contain RNA genomes [7, 8], indicating that RNA viruses are as successful in rice as in other plants. As shown in dicot plants, the RNA silencing pathway also mediates a potent antiviral defense against RNA viruses in rice by producing 21- and 22-nucleotide (nt) virus-derived small interfering RNAs (siRNAs) to confer antiviral RNA interference (RNAi) [4, 9–11].

Several properties of rice viruses may explain why viral diseases remain a constant threat to rice production. Because of their error-prone replication mechanisms, RNA viruses possess extraordinary adaptive abilities to escape effector-triggered immunity (ETI) that depends on specific recognition of pathogen proteins by disease resistance genes. The vast majority of rice viruses are transmitted in field by insect vectors such as leafhoppers and planthoppers [7, 8]. Surprisingly, although aphid-transmitted viruses such as cucumoviruses and potyviruses are widespread in crops, no sequenced rice virus has been shown to spread by aphid vectors [10, 12, 13]. Notably, insect-borne viruses emerge and re-emerge to cause devastating rice disease epidemics without clearly defined origins. These include rice tungro viruses, rice grassy stunt virus, rice ragged stunt virus and rice gall dwarf virus emerged in 1960’s and 1970’s as well as Southern rice black-streaked dwarf virus (SRBSDV) that began to circulate over two decades ago [14–19]. Another intriguing property of some rice insect-borne viruses is that they can suddenly become widespread after several years of little or no infection [7, 8].

It is well known that some of the most dangerous human viruses have zoonotic origins from wild animals and/or depend on animals as the reservoir or intermediate hosts [20–24]. There is evidence for weed species to act as the reservoir hosts for rice viruses. For example, both rice tungro spherical virus (RTSV) and rice tungro bacilliform virus (RTBV) have been detected in a wide range of weed plants although leafhopper recovery of virus from infected weeds to rice has not been convincingly established [14]. Importantly, it remains unknown whether native plants host new endemic viruses that can spread to cause diseases in cultivated rice varieties, thereby acting as emerging pathogens of rice.

Recent technological advances have dramatically accelerated the pace of RNA virus discovery both on land and in the ocean [25–28]. One metagenomics approach that has been most successful in plants is virus discovery by deep sequencing and assembly of virus-derived siRNAs (vdSAR), which is based on the findings that the viral siRNAs produced by plant and invertebrate hosts overlap in sequence and can be assembled into the viral genome segments for virus discovery [29–32].

In this work, we use vdSAR to identify new viruses of native plants with spillover potential to cause disease epidemics of rice. We discover a new positive-strand RNA virus from colonies of Asian wild rice (*O. rufipogon*) in a genetic reserve and demonstrate virus recovery from infected wild rice plants to *O. sativa* seedlings by one specific aphid species. We further show that a molecular clone of the identified virus establishes systemic infection and causes significantly reduced tillering in a popular rice cultivar. Our results indicate that novel rice viral pathogens may emerge from native habitats, a rare aphid transmitted viral pathogen poses serious threat to rice cultivar.

## Results

### Discovery of a new positive-strand RNA virus from wild rice colonies

We hypothesized that wild native plants related to rice species (*Oryza sativa*) host endemic viruses pathogenic to the domesticated rice cultivars. Guangxi University is home to a genetic reserve of wild rice colonies [33] collected across China and vegetatively maintained in net-houses since 1999. We thus sequenced pools of small RNAs from ~1000 individual wild rice colonies to search for new viruses using vdSAR as described previously [30]. vdSAR identified contigs of assembled small RNAs from one pool of Asian wild rice (*O. rufipogon*) colonies (Figs 1A-1B and S1) that exhibit strong sequence similarity to sugarcane yellow leaf virus (ScYLV) from the positive-strand RNA virus genus *Polerovirus* [34]. RT-qPCR detection of the coat protein (CP) gene (Fig 1B) from the identified virus, designated subsequently rice tiller inhibition virus (RTIV) according to the disease symptom induced in domesticated rice plants (see below), indicated that 6 of the 10 colonies in the pool were infected with RTIV (S2 Fig).

**Fig. 1.**
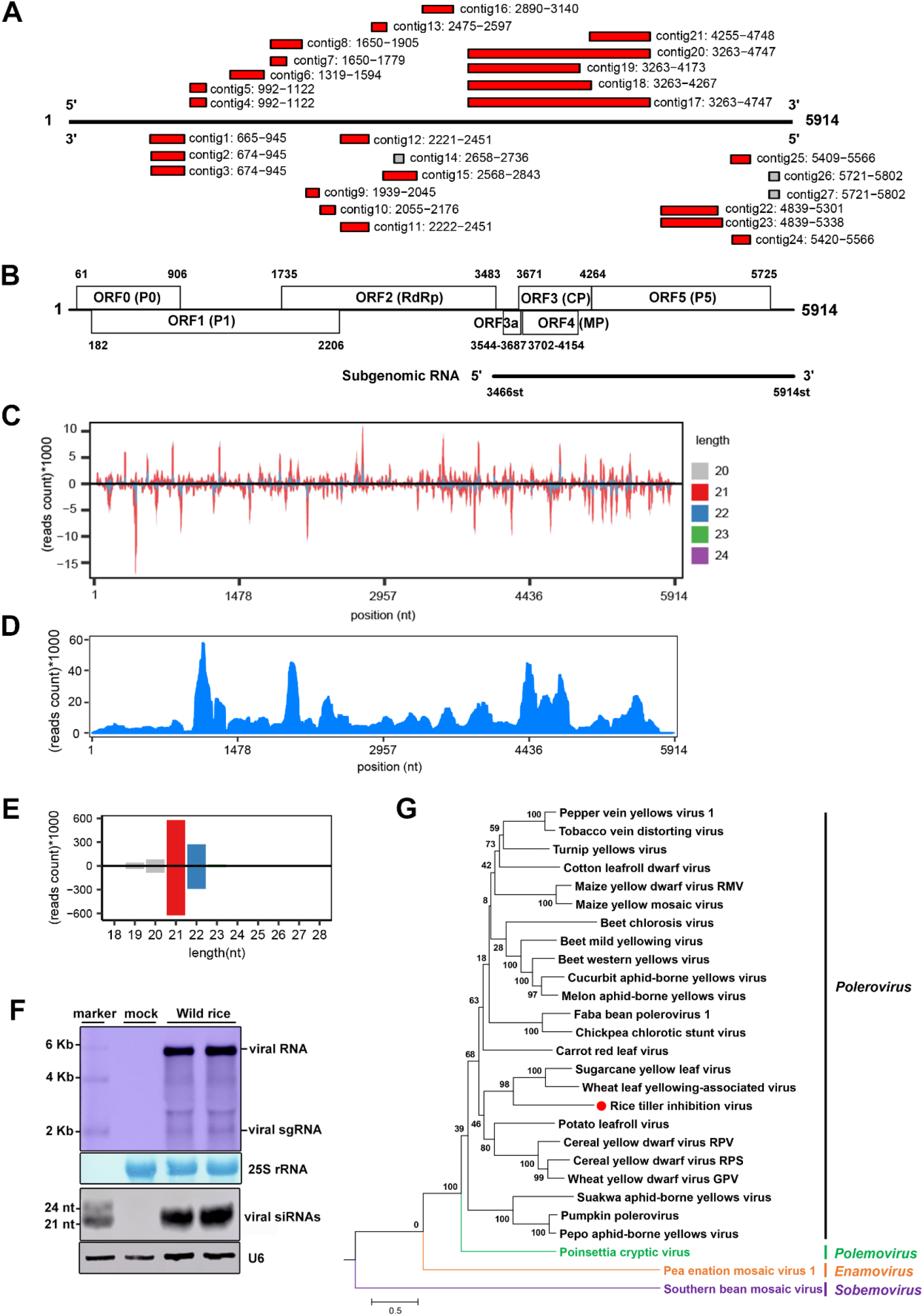
The discovery of RTIV in Asian wild rice colonies. **(A)** Assembled RTIV contigs from vdSAR. **(B)** Schematic representation of the RTIV genome and subgenomic RNA, with the six main ORFs (ORF 0-5) and a small ORF3a. **(C)** Small RNA-Seq of Wild-rice individual, the single-nucleotide resolution maps of the positive- and negative-strand 20-24nt vsiRNAs in the viral genome from the sequenced small RNA pool. **(D)** The RTIV genome maps of RNA-seq reads sequenced from colony no. 9 wild rice plants. **(E)** Size distribution of the positive- and negative-strand small RNA reads derived from RTIV. **(F)** Northern blot detection of the viral genomic/subgenomic (sg) RNAs (top 2 panels) and vsiRNAs (bottom 2 panels) in two individual wild rice plants from colony no. 9. 25S rRNA was stained and U6 was probed to show equal loading. **(G)** Phylogenetic relationship of RTIV with known members of *Polerovirus* and other genera in the family *Solemoviridae* based on the RdRP.

RTIV was confirmed using vdSAR by small RNA sequencing of colony no.9 (Figs 1C and S3). The full genomic sequence of RTIV was then obtained by sequencing the total RNAs depleted of the ribosomal RNAs from colony no. 9 (Fig 1D) and by rapid-amplification of cDNA ends (RACE) for determining the viral 5’ and 3’ terminal sequences (Figs 4B, S4A and S4C). Results from these analyses showed that RTIV genome was 5,914 nucleotides (nt) in length (Table S1) and encoded seven open reading frames (ORFs) as found in known poleroviruses [35, 36] (Fig 1B). The sequence of 5’-terminal six nucleotides of the RTIV genomic RNA (5’- ACAAAA-3’) was identical to that of the known poleroviruses (S4B Fig). Further phylogenetic analysis of the viral RdRPs indeed showed that RTIV is closely related to wheat leaf yellowing-associated virus (WLYaV) and ScYLV within poleroviruses in the *Solemoviridae* family [36] (Fig 1G). Homologic identities of each RTIV individual proteins or its genomic nucleotide sequences to known poleroviruses shows that RTIV and WLYaV CPs exhibited 80.1% sequence identity, which is the highest detected between an RTIV-encoded protein and known viral proteins (Table 1). Species demarcation criteria in the genus *Polerovirus* is a >10% difference in amino acid sequence identity of any gene product from its closest relative [35]. Thus, we propose naming RTIV as a new species in the *Polerovirus*.

**Table 1.** Identity percentage of viral proteins and nucleotide sequences of RTIV with known members in Polerovirus. Identity percentage of amino acid and nucleotide was analyzed by Sequence Demarcation Tool. Accession number of known poleroviruses used in the analysis was listed in Table S2.

| RTIV<br>*Polerovirus* | ORF0<br>(aa) | ORF1<br>(aa) | ORF1-2<br>fusion<br>(aa) | ORF3<br>(aa) | ORF3a<br>(aa) | ORF4<br>(aa) | ORF3-5<br>fusion<br>(aa) | Full<br>length<br>(nt) | 5'UTR<br>(nt) | 3'UTR<br>(nt) |
| --- | --- | --- | --- | --- | --- | --- | --- | --- | --- | --- |
| Wheat leaf yellowing-associated virus | 20.7 | 34.8 | 48.9 | 80.1 | 63.6 | 67.3 | 62.7 | 64.5 | 55.2 | 52.0 |
| Sugarcane yellow leaf virus | 23.5 | 33.6 | 48.7 | 77.6 | 86.4 | 66.0 | 62.6 | 65.1 | 62.7 | 59.4 |
| Potato leafroll virus | 24.2 | 34.0 | 46.8 | 42.1 | 52.3 | 30.5 | 33.1 | 60.1 | 66.7 | 55.7 |
| Pepper vein yellows virus 1 | 23.3 | 35.2 | 46.1 | 48.0 | 61.4 | 27.2 | 37.5 | 60.9 | 66.0 | 66.3 |
| Maize yellow mosaic virus | 23.3 | 34.4 | 46.1 | 41.8 | 61.4 | 28.8 | 43.5 | 59.6 | 62.8 | 57.2 |
| Maize yellow dwarf virus RMV | 23.5 | 32.9 | 45.8 | 41.3 | 34.1 | 32.9 | 44.3 | 60.3 | 55.6 | 65.8 |
| Turnip yellows virus | 25.2 | 34.7 | 45.7 | 44.6 | 52.3 | 31.3 | 33.1 | 58.7 | 72.4 | 54.7 |
| Faba bean polerovirus 1 | 21.6 | 31.3 | 45.4 | 44.6 | 56.8 | 35.1 | 33.2 | 60.2 | 85.2 | 55.0 |
| Chickpea chlorotic stunt virus | 23.3 | 32.4 | 45.4 | 41.8 | 52.3 | 30.0 | 35.9 | 59.8 | 63.8 | 57.1 |
| Carrot red leaf virus | 18.0 | 30.7 | 45.3 | 42.3 | 50.0 | 31.5 | 34.3 | 59.3 | 63.0 | 61.4 |
| Tobacco vein distorting virus | 25.0 | 32.7 | 45.2 | 47.7 | 59.1 | 29.5 | 32.6 | 59.3 | 60.4 | 58.7 |
| Beet mild yellowing virus | 22.0 | 32.9 | 44.5 | 42.3 | 42.6 | 32.4 | 32.9 | 59.9 | 61.3 | 58.7 |
| Cucurbit aphid-borne yellows virus | 19.9 | 31.7 | 44.2 | 46.1 | 56.8 | 27.3 | 38.3 | 58.8 | 84.2 | 51.9 |
| Wheat yellow dwarf virus GPV | 22.0 | 30.9 | 44.2 | 44.6 | 54.5 | 30.9 | 33.2 | 60.0 | 61.7 | 60.2 |
| Melon aphid-borne yellows virus | 21.9 | 30.3 | 44.2 | 42.3 | 51.1 | 30.9 | 36.2 | 59.6 | 84.2 | 59.9 |
| Beet western yellows virus | 23.1 | 32.6 | 43.9 | 45.1 | 36.4 | 33.1 | 33.3 | 59.8 | 67.9 | 69.3 |
| Cotton leafroll dwarf virus | 20.5 | 29.8 | 43.5 | 49.2 | 56.8 | 33.3 | 38.8 | 59.5 | 74.5 | 55.2 |
| Cereal yellow dwarf virus RPV | 21.6 | 30.3 | 43.3 | 46.9 | 52.3 | 28.4 | 38.0 | 60.5 | 68.3 | 53.5 |
| Cereal yellow dwarf virus RPS | 23.1 | 29.7 | 43.2 | 45.4 | 50.0 | 33.1 | 37.9 | 61.1 | 63.0 | 57.9 |
| Pepo aphid-borne yellows virus | 21.3 | 29.5 | 42.6 | 46.1 | 56.8 | 30.0 | 37.6 | 59.1 | 62.5 | 57.5 |
| Pumpkin polerovirus | 20.2 | 28.5 | 42.2 | 45.6 | 56.8 | 27.3 | 36.7 | 59.7 | 62.5 | 52.9 |
| Beet chlorosis virus | 20.3 | 26.6 | 41.6 | 46.1 | 54.8 | 31.8 | 32.4 | 59.4 | 72.0 | 50.6 |
| Suakwa aphid-borne yellows virus | 23.5 | 26.6 | 40.6 | 46.6 | 48.9 | 31.8 | 37.6 | 59.0 | 70.5 | 56.2 |

The total small RNA reads from colony no. 9 mapped to the full-length genome of RTIV displayed strong size preference for 21- and 22-nt with a minor peak at 20-nt (Fig 1C and 1E). This size distribution pattern is typical of those shown previously for virus-derived siRNAs (vsiRNAs) produced by plant antiviral RNAi response to RNA virus infection [9]. Moreover, all three peaks of RTIV vsiRNAs exhibited approximately equal ratios of the positive and negative strands (Fig 1C and 1E) and the viral full-genomic RNA was targeted by the positive- and negative-strand vsiRNAs. These properties of RTIV vsiRNAs are consistent with the model of plant vsiRNA biogenesis [9] in which viral long dsRNA precursors are processed into 21- and 22-nt vsiRNAs by Dicer-like 4 (DCL4) and DCL2, respectively. Notably, millions of RNA-seq reads of colony no.9 can also be mapped to the full-length genome of RTIV (Fig 1D), indicating its successful adaptation and propagation in the native habitat.

Total RNAs were also extracted from two individual plants vegetatively propagated from no. 9 colony for Northern blot detection of RTIV high and low molecular weight RNAs. RTIV vsiRNAs in the size range of 21- to 22-nt accumulated to readily detectable levels in both of the wild rice plants (Fig 1F). The largest and the most abundant RNA molecule detected in both plants exhibited a size approximately 6 kilobases (kb) in length and thus most likely corresponded to the full-length genomic RNA of RTIV (Fig 1F). A smaller viral RNA species detected in the infected plants was identified as the 2,449-nt polerovirus-specific subgenomic RNA (sgRNA) for CP expression by 5’ RACE and Northern blot hybridization with region specific probes [37–39] (Figs 1B, 1F and S5). These results further demonstrate induction of a typical antiviral RNAi response and production of both the viral genomic and subgenomic RNAs in individual wild rice plants, indicating an active RTIV infection of the Asian wild rice colony.

Together, our findings discovered a new polerovirus that has established persistent infection in wild rice colonies maintained vegetatively for decades in the net house.

### Recovery of RTIV from persistently infected wild rice plants by vector aphids for successful transmission and infection of a cultivated rice species

We next investigated whether RTIV detected in wild rice colonies remains infectious and can spread to cause infection in domesticated rice species (*O. sativa*). Poleroviruses spread in nature exclusively by specific aphid vectors in a persistent non-propagative manner [7, 8, 36, 40]. Thus, we attempted to recover RTIV from infected Asian wild rice plants by four aphid species known to transmit related viruses in cereals: *Rhopalosiphum padi (R. padi)*, *Schizaphis graminum*, *Sitobion avenae* and *Metopolophium dirhodum* [36, 41, 42]. Aphids were fed on the RTIV-positive wild rice plants for 3 days before they were transferred to seedlings of wildtype *O. sativa* spp. *Japonica Nipponbare* (*NIP*) or knockout for the *Argonaute 18* gene (*AGO18*) which enhances the expression of AGO1, the essential antiviral RNAi effector protein [43].

We found that both NIP and *ago18* mutant plants developed a reduced tillering symptom after inoculation only with R. padi fed on the RTIV-positive wild rice (Fig. 2A). Northern blotting detected the modestly higher accumulation of the viral genomic and subgenomic RNAs as well as the vsiRNAs in the symptomatic *ago18* mutants compared to NIP plants (Fig. 2B), probably because of the compromised antiviral RNAi in *ago18* plants. Using the antisera raised against CP of RTIV, we also detected comparable expression of the viral CP with the predicted 22 kD in these symptomatic plants by Western blotting (Fig. 2B). These findings indicated that R. padi aphids recovered RTIV from the infected wild rice plants and transmitted the virus to the cultivated rice species for infection and disease induction, though antiviral RNAi was induced to prevent its infection in plants.

**Fig. 2.**
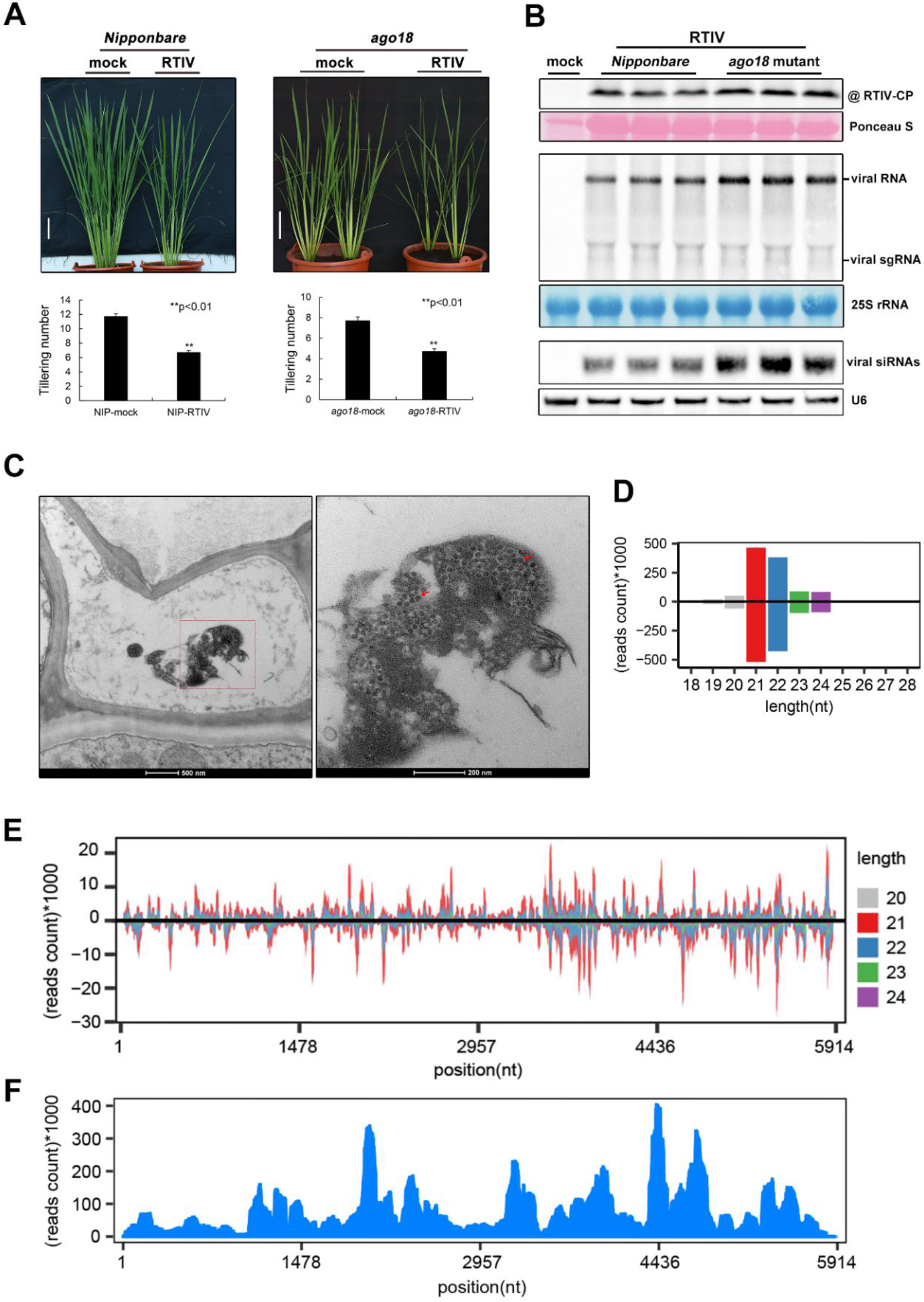
Recovery of RTIV from wild rice and transmission to the domesticated rice by aphid *R. padi*. **(A)** The reduced tillering symptom of *ago18* plant (A) or *NIP* plant photographed at ~40dpi. Statistic analysis of tiller number for RTIV-infected *ago18* plants or *NIP* plants compared to mock plants, 25 plants used for each analysis. Scale bars =10 cm. **(B)** Detection of RTIV CP by Western blotting (top panel), RTIV genomic/subgenomic (sg) RNAs (middle panel) and RTIV siRNAs (bottom panel) by Northern blotting in *Nipponbare* and *ago18* mutant plants. Loading control was monitored by ponceau staining, 25S rRNA staining or U6 RNA probing in the same membrane. **(C)** Distribution of RTIV virions within the cell of *ago18* mutant plants with a magnified view of the boxed area shown at the right. Viral particles (red arrows) are approximately 30 nm in diameter. **(D/E)** Size distribution (D) and the single-nucleotide resolution maps (E) of the positive- and negative-strand RTIV-derived small RNA reads (18-30 nt) sequenced from RTIV-infected *ago18* plants after aphid transmission. **(F)** The RTIV genome maps of RNA-seq reads sequenced from RTIV-infected *ago18* plants after aphid transmission.

We then sequenced the total RNA and small RNAs in the symptomatic *ago18* plants after inoculation with viruliferous *R. padi* as described above for wild rice plants. The results from the assembly of the RNA or small RNA reads demonstrated the infection of the symptomatic *ago18* plants with RTIV (Fig 2E and 2F). Both the size distribution and strand ratio of the RTIV vsiRNAs from the RTIV-infected *ago18* plants were indistinguishable from those of the RTIV-infected wild rice plants (Fig 2D and 2E). Notably, no contigs assembled from the large or small RNA reads shared significant sequence similarities with any of the known viruses. Moreover, observation with transmission electron microscope (TEM) found large quantity of spherical viral particles approximately 30 nm in diameter in the vascular bundle of the symptomatic *ago18* plants (Fig 2C). Therefore, we conclude that the Asian wild rice colonies in the genetic reserve contain infectious RTIV particles that remain competent not only in independent vector transmission by *R. padi*, but also in the systemic infection of a cultivated rice species.

### Independent infection and disease induction of wild type cultivated rice plants by cloned RTIV

We further developed the infectious molecular clone of RTIV to verify its infectivity and pathogenesis. A binary plasmid (pCass-RZ-RTIV) containing the full-length cDNA of RTIV RNA genome was cloned between the 35S promoter and a ribozyme in a plasmid vector modified from pCass reported previously [50, 51] (S6 Fig). To test the infectivity of pCass-RZ-RTIV, leaves of both wild type and *DCL2*- and *DCL4*-RNAi knockdown *Nicotiana benthamiana* plants (*dcl2/4-RNAi*) were firstly inoculated by infiltration with *Agrobacterium tumefaciens* that carried pCass-RZ-RTIV. Northern blot hybridization detected accumulation of the viral genomic RNA and the viral subgenomic RNA as well as the 24nt vsiRNAs in the infiltrated leaves of *N. benthamiana dcl2/4-RNAi* plants (Fig 3A, right 3 lanes). By comparison, much lower levels of the viral genomic and subgenomic RNAs and vsiRNAs were detected in the infiltrated leaves of wild-type *N. benthamiana* plants (Fig 3A, left 3 lanes), which is consistent with reduced accumulation of RTIV in NIP compared to *ago18*. These results suggest that the cloned RTIV genome initiates replication in the infiltrated leaves and directs synthesis of the viral subgenomic RNA, and triggers the biogenesis of the vsiRNAs which could curb RTIV propagation.

**Fig. 3.**
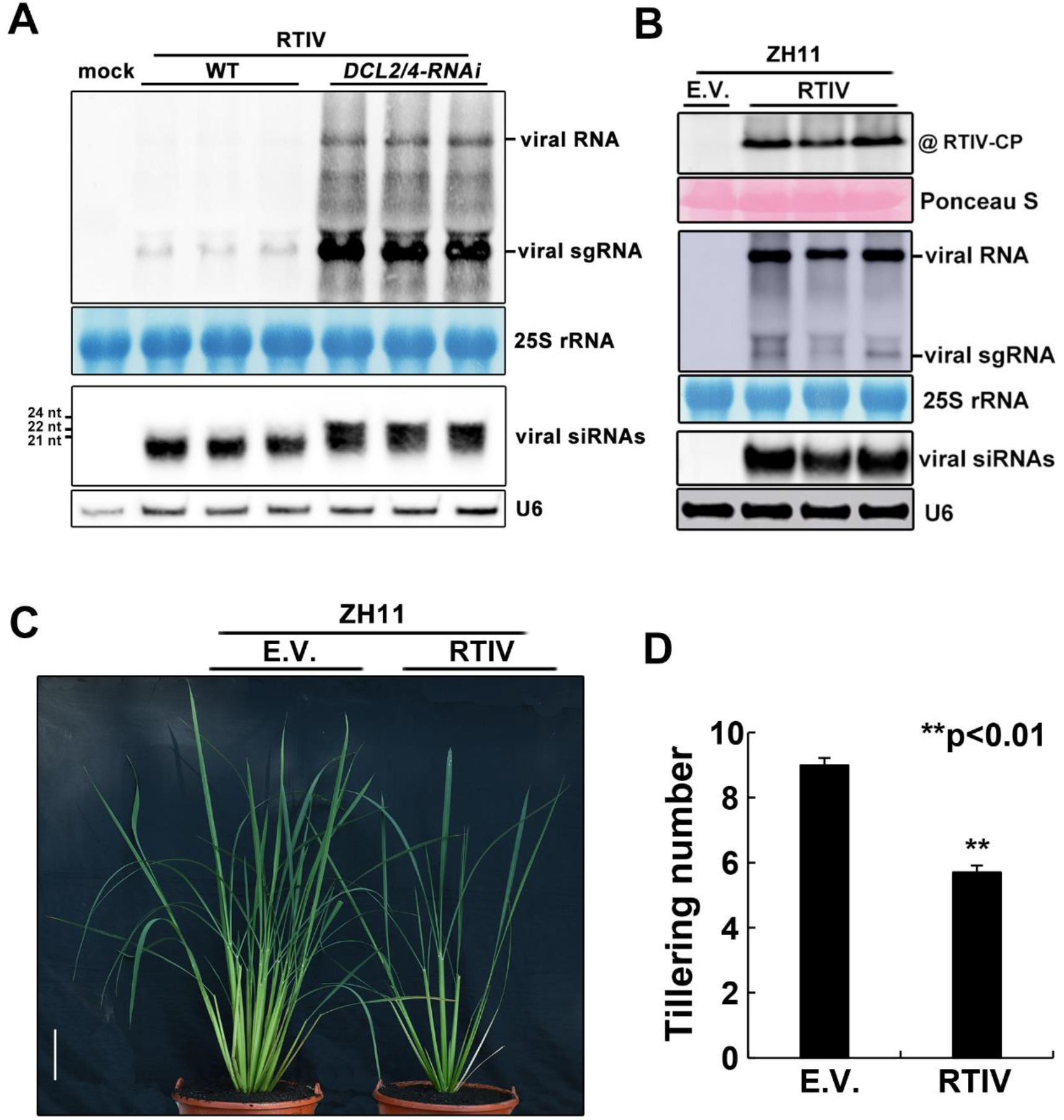
Infection and disease induction of an elite rice variety by cloned RTIV. **(A)** Northern blot detection of the viral genomic/subgenomic RNAs and the vsiRNAs in the leaves of wild type and DCL2/4-knockdown *N. benthamiana* plants infiltrated with pRTIV-Agrobacteria. 25S rRNA was stained or U6 RNA probed in the same membrane to show equal loading. **(B)** Detection of RTIV CP by Western blotting (top panel), RTIV genomic/subgenomic (sg) RNAs (middle panel) and RTIV siRNAs (bottom panel) by Northern blotting in upper uninoculated leaves of Zhonghua-11 rice plants after Agro-inoculation. Loading control was monitored by ponceau staining, 25S rRNA staining or U6 RNA probing in the same membrane. **(C)** The reduced tillering symptom of Zhonghua-11 plants ~6 weeks after Agro-inoculation with RTIV. **(D)** Statistical analysis of tiller numbers of RTIV-infected Zhonghua-11 or mock (n=30), \*\**p*<0.01 determined using the Student’s test and mean significant difference. Error bar: standard deviations, Scale bars: represent 10 cm.

The variation of RTIV accumulation in wildtype and antiviral RNAi-defective plants indicates that RTIV may not possess potent VSR to suppress antiviral RNAi in plants. P0 protein is VSR of polerovirus functioning to mediate AGO1 degradation [52–53]. We found that the annotated P0 protein of RTIV showed very low identity (~20%) to known P0 of other poleroviruses (Table 1). When tested, it did not show VSR activity in transient expression-induced gene silencing in 16c GFP transgenic plants which is routinely used for examining VSR activity [54] (S7 Fig). Coat Protein (CP), Movement protein (MP) or full length of RTIV did not exhibit VSR activity either when tested in 16c plants (S7 Fig). In addition, Western blotting results showed that the expression level of AGO1a protein in RTIV-infected rice plants was not decreased compared to mock rice plants (S8 Fig). These results suggest RTIV is a specific VSR-lacking polerovirus infecting rice.

Next, we used an Agro-inoculation procedure developed for rice seedlings [55] to determine whether the cloned RTIV can infect wild type Zhonghua-11 plants, an elite Chinese variety of Asian cultivated rice (*O. sativa* spp. *Japonica*). We found that the viral genomic and subgenomic RNAs, the vsiRNAs as well as CP were all readily detectable in the non-infiltrated leaves of independently inoculated plants 6 weeks post-inoculation (Fig 4B), indicating systemic infection of wild-type rice plants by RTIV. RTIV-infected Zhonghua-11 plants also exhibited the symptom of significantly reduced tillering, an important trait of rice yield [56], compared to mock- and RTIV-infected Zhonghua-11 plants (Fig 3C, 3D). However, RTIV-infected rice plants did not display additional symptoms such as dwarf or color distortion on leaves frequently associated with the increased or decreased tillering symptom of rice viral diseases [7, 8]. These findings demonstrate that the infectious clone of RTIV independently establishes systemic infection in the Asian cultivated rice species and causes tiller inhibition, which further indicates that native plants host an endemic virus with spillover potential as a new pathogen into cultivated rice (*Oryza sativa*).

**Fig. 4.**
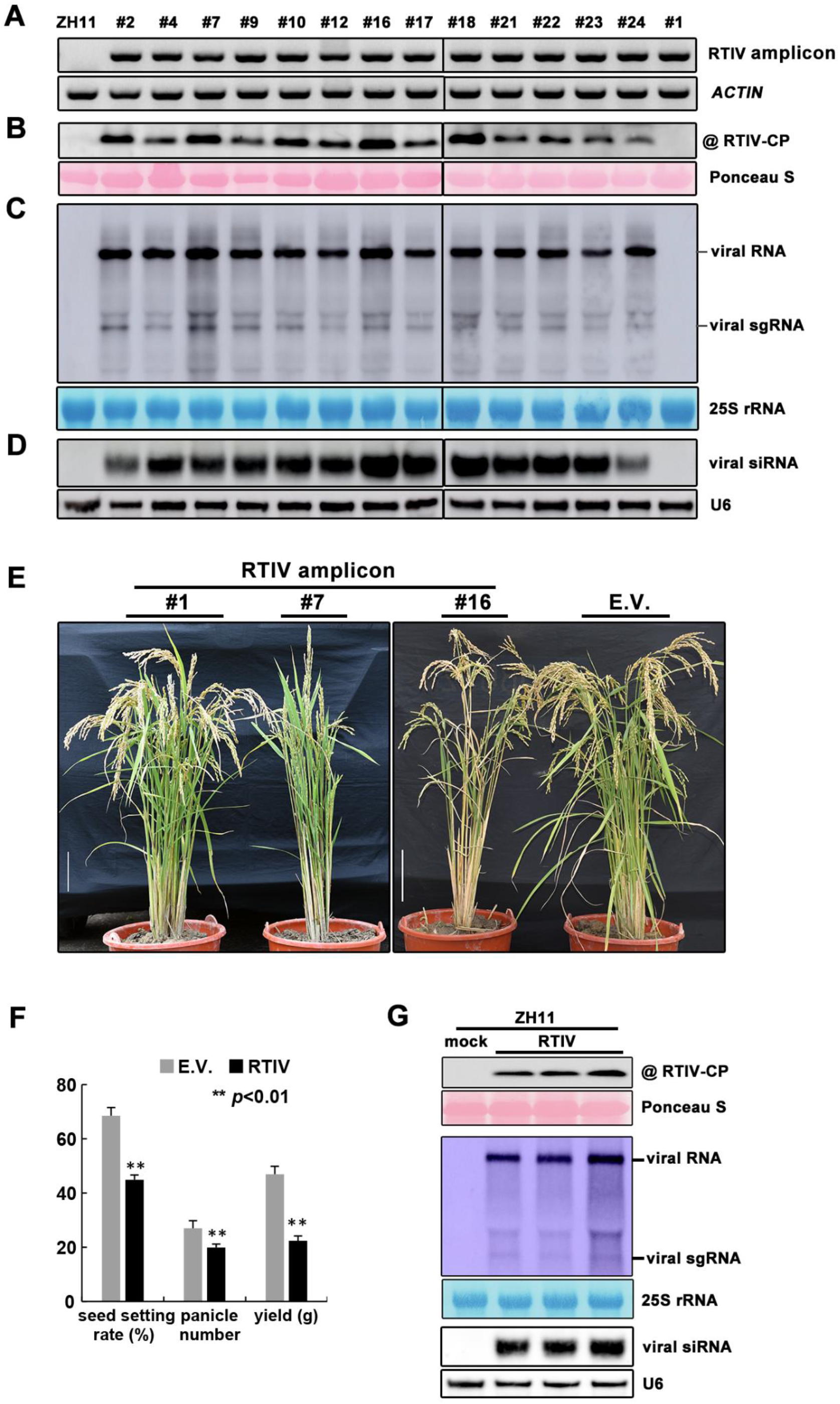
RTIV replication and pathogenesis in rice cultivar induced by RTIV encoded in a stably integrated transgene. **(A)** PCR detection of the RTIV transgene in transgenic Zhonghua-11 plants. (b-d) Detection of RTIV CP by Western blotting. **(B)** RTIV genomic/subgenomic (sg) RNAs **(C)** and RTIV siRNAs **(D)** by Northern blotting in independent lines of Zhonghua-11 rice plants carrying a stable virus transgene. Loading control was monitored by ponceau staining, 25S rRNA staining or U6 RNA probing in the same membrane. **(E)** Phenotypes of transgenic Zhonghua-11 plants carrying an empty vector transgene (E.V.), a non-replicating (#1) or replication-competent (#7 and #16) RTIV transgene. Scale bars =10 cm. **(F)** The seeds set rate, panicle number and individual plant yield of E.V. and RTIV replication-competent transgenic rice. \*\**p*<0.01 determined using the Student’s test and mean significant difference. Error bar: standard deviations. **(G)** Detection of RTIV CP, genomic/subgenomic (sg) RNAs and vsiRNAs by Western blotting or Northern blotting in healthy Zhonghua-11 rice plants 2 weeks after RTIV transmission from above RTIV-positive transgenic rice plants, loading control was monitored by ponceau staining, 25S rRNA staining or U6 RNA probing in the same membrane.

### Replication and disease induction by RTIV encoded in a stably integrated transgene

We took an independent approach to examine the infectivity of the cloned RTIV in Zhonghua-11 plants by stable plant transformation. We obtained 14 lines of stable transgenic Zhonghua-11 plants carrying an RTIV transgene driven by the 35S promoter. Active RTIV replication was documented in 13 of these transgenic lines by Northern blot detection of the viral genomic and subgenomic RNAs as well as the vsiRNAs and by Western blot detection of CP expression (Figs 4A-4D and S9). Notably, all of the 13 lines of RTIV replication-competent transgenic plants developed the severe disease symptom of significantly reduced tillering compared to the control plants carrying either the empty vector or a non-replicating RTIV transgene (Fig 4E), and the accumulation of viral CP protein, viral RNAs and vsiRNA was even higher compared to that in wild rice plants, *NIP* infected RTIV through aphid or ZH11 mechanically infected RTIV molecular clone (S9 Fig). Further statistical analyses showed that panicle number, seed setting or yield of single plant were all significantly reduced in the transgenic plants supporting active RTIV replication, compared to the control plants carrying an empty vector transgene (E.V.) (Fig 4F).

We further examined whether RTIV can be transmitted from this transgenic rice cultivar by its aphid vector. Aphids *R. padi* were fed on the RTIV-positive transgenic rice plants for 3 days and then transferred to healthy seedlings of wildtype Zhonghua-11 plants. After 2 weeks, viral CP, RNA genomic and subgenomic RNAs, and vsiRNAs were detected in these wildtype rice plants by Western blot or Northern blot respectively (Fig 4G). Consistently, low-tillering disease was also developed in these rice plants after 45 dpi (S10 Fig). These results further confirm that RTIV newly discovered from Asian wild rice is an aphid-transmitted viral pathogen of rice plants and capable of causing severe disease to threaten cultivated rice.

### Altered expression pattern of defense and tillering-relevant genes in rice cultivar after RTIV infection

The severe disease induction in rice cultivar suggested abnormal homeostasis in plants caused by RTIV infection. With attempt to find out the disturbed pathways which could involve in the disease induction, we analyzed the global transcriptome based on the total RNA data previously sequenced in symptomatic *ago18* plants. We found that expression of over thousand genes was strikingly altered in the diseased rice plants (Fig 5A). Among hundreds of upregulated genes, most of them have been reported to regulate stress response, defense response, or responding to salicylic acid, indicating the induced antiviral response after viral infection (Fig 5D) [44, 45]. On the contrary, among down-regulated genes, most of them are related to the biosynthesis of secondary metabolites (Fig 5B), suggesting an adverse effect on normal growth and development of plants. Notably, we found that expressions of several tillering-relevant genes including CCD8B, D53, ERG1 (Rpp17) and MOC1 were significantly increased by more than 2 times in the symptomatic plants compared to mock plants (Fig 3C) [46–49]. We further confirmed the expression alternation of these specific genes by RT-qPCR analysis (Fig 5E). Therefore, these results indicate the novel rice viral pathogen may cause viral disease by disturbing the defense and tillering pathways of rice plants.

**Fig. 5.**
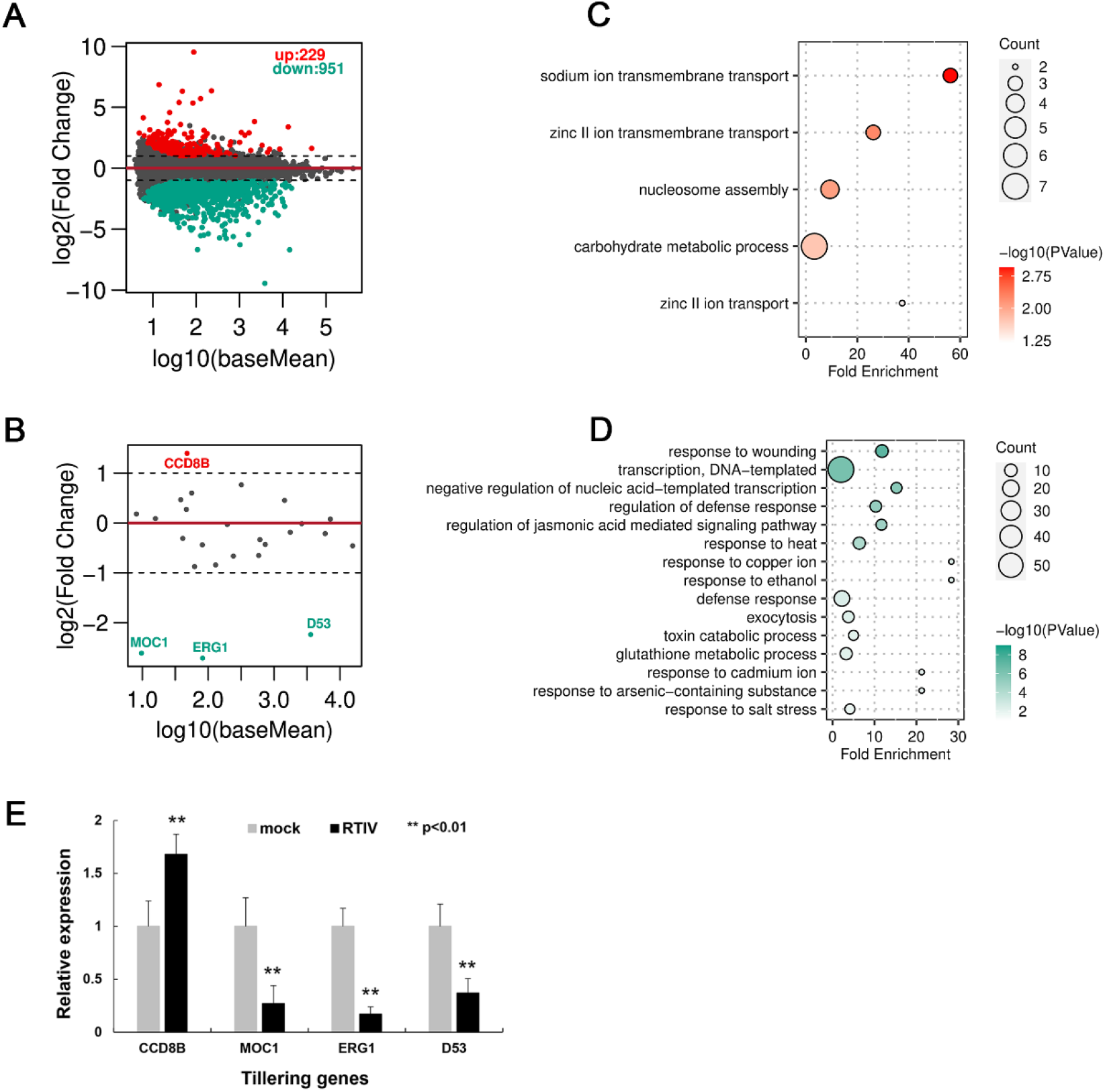
RTIV modulated defense responses and caused low-tillering disease in rice plants after spillover through vector aphid. **(A)** All genes of altered expression profile, log2 values are the fold change of gene expressions, baseMean are the gene reads of library standardization. **(B)** Tillering-related genes of altered expression profile. **(C-D)** KEGG pathway enrichment analysis of up-regulated genes (C) or down-regulated genes (D), count indicates the number of gene, log10 (*p* value) indicates significance of difference. **(E)** RT-qPCR results shown transcript level of tiller genes in *ago18* plants after RTIV infection compared to non-infected mock plants. Relative expression level of each gene was normalized to reference gene *OsActin*. Experiment was repeated 3 times, with 3 replicates for each reaction. \*\**p*<0.01 determined using the Student’s test and mean significant difference. Error bar: standard deviations.

## Discussion

An endemic virus of native plants has spillover potential to cause disease epidemics in crops, when the virus can establish virulent infection in crops and can spread from infected native plants to crops by insect vectors or other efficient mode of transmission. Much is known about the host range of rice viral pathogens, which helps to identify the reservoir weed plants and the source of inoculum in new season of rice cultivation [7, 8]. Use of small RNA sequencing in this work led to the discovery of RTIV that establishes persistent infection in Asian wild rice colonies and has spillover potential as an emerging pathogen of rice.

In the wild rice colonies maintained vegetatively for decades, we detected accumulation of the genomic and subgenomic RNAs of RTIV and induction of the antiviral RNAi response, indicating long-term persistent infection in these native plants. We show that Agro-inoculation with a molecular clone of RTIV induces systemic infection and causes a unique low-tillering disease in an elite variety of Asian cultivated rice species. Moreover, stable transgenic lines of the rice variety that carries an RTIV transgene support robust virus replication and develop the low-tillering disease. Potato leafroll virus (PLRV, a *Polerovirus* as RTIV) also replicates efficiently in tobacco and potato plants stably transformed with a PLRV transgene and transgenic lines of potato develop severe disease symptoms [57]. Omura and colleagues have shown that persistent rice dwarf virus infection of Asian rice plants maintained vegetatively for six years triggers nonsense mutations in the dsRNA genome of the reovirus and complete loss of transmissibility by its leafhopper vectors [58]. Notably, we demonstrate recovery of RTIV by *R. padi* aphids and successful transmission to rice seedlings from the persistently infected wild rice plants. Together, our findings provide the first evidence to support native plants as a possible origin of emerging rice viral pathogens.

Aphid-transmitted rice viral pathogens have been rarely reported. The first determined aphid-borne rice virus and its aphid vector *R. Padi* in this work will facilitate future study in the field and facilitate specific monitoring of the virus in natural fields. Notably, *R. padi* is a global pest and the vector of diverse yellow dwarf viruses (YDVs) of cereals from the genera *Polerovirus* and *Luteovirus* of the *Solemoviridae* and the *Tombusviridae*, respectively [35, 36, 59, 60], and Rice infection with undefined YDVs such as rice giallume virus have been found in Europe [8, 61]. Future metagenomic sequencing analysis will find whether RTIV-related viruses spread worldwide to threaten cultivated rice, and whether more novel viruses with potential threat are widespread in native plants.

## Materials and Methods

### Plant material and insect vectors

Wild rice plants were maintained in the outdoor genetic reserve net-houses at Guangxi University, China as described previously[33]. *Nicotiana benthamiana* plants were cultivated in a growth chamber under 16 hr light/8 hr dark at 24°C. *N. benthamiana-DCL2/4-RNAi* [62] were kindly provided by Dr. A.L.N. Rao at the University of California-Riverside. Zhonghua-11 rice plants (*Oryza sativa*) were grown in a greenhouse maintained at 26 °C with a 14/10 hr (light/dark) photoperiod and 70% relative humidity. Rice *ago18* mutant was described previously [43]. Aphid vectors were reared on rice seedlings in clear containers in a controlled environment at 23°C [63].

### Small RNA sequencing, analysis of siRNAs and contig assembly by vdSAR

Rice RNA samples were sent to Beijing Genome Institute (BGI, Shenzhen, China) for deep sequencing of small RNAs by MGISeq-2000 platform. In brief, small RNAs were ligated with RNA adaptors and reversely transcribed into cDNAs that were amplified by PCR and subjected to sequencing. After sequencing, adapter sequences were trimmed and the reads in 18~28nt size range were retained by cutadapt software.

Clean reads were processed by Velvet [64] for de novo assembly. Contigs ≥100nt were subjected to local BLAST for BLASTn and BLASTx against nt and nr database respectively. The contigs with the best match to viral protein or nucleotide sequence and E-values <1×10−3 were identified as viral contigs. Viral contigs were further verified and assembled manually in Seqman program. Bowtie software [65] was used for obtaining vsiRNA by mapping sRNA reads to viral contigs and RTIV genome with perfect alignment and two mismatches, respectively [30]. The number of vsiRNAs mapped to RTIV genome were calculated and visualized by R scripts.

### RNA sequencing and analysis

Rice RNA samples were also sent to BGI for RNA deep-sequencing after removing rRNAs by MGISeq-2000 platform. Briefly, after rRNAs removal, total RNAs were ligated with RNA adaptors and reversely transcribed into cDNAs for further amplification by PCR and sequencing. From the generated data, the RNA fragments were mapped to RTIV genome by using bowtie2 and visualized with R program.

### RNA extraction, Northern blot analysis

High and low molecular weight RNAs were isolated by hot phenol method. Northern blotting was conducted as our previously reported method [66] with biotin-labeled DNA probes using Biotin-11-dUPT (Thermo Scientific USA). High-molecular weight RNAs were separated in 1 % formaldehyde agarose gels and transferred to Hybond-N^+^ membranes (GE Healthcare, UK) for RNA hybridization in membranes. Low-molecular weight RNAs were separated in 15 % urea denaturing gels and transferred to Hybond-NX membranes (GE Healthcare, UK) for hybridization with biotin-labeled DNA oligo probes and detectiond by Chemiluminescent Nucleic Acid Detection Module Kit (Thermo Scientific, USA).

### Phylogenetic analysis and sequences identities analysis

Sequences of other viruses in the family of Solemoviridae and Tombusviridae (Table S2) were retrieved from NCBI (http://www.ncbi.nlm.nih.gov/). The tree was assembled using MEGA 5.22 and the maximum-likelihood method based on the WAG + G+I +F model. Numbers on the nodes indicate the percentage of bootstrap replicates supporting the branch (n=1000). Percent sequence identities of amino acid and nucleotide were analyzed by Sequence Demarcation Tool.

### Insect transmission

Virus-free aphids *Rhopalosiphum padi*, *Schizaphis graminum*, *Sitobion avenae* and *Metopolophium dirhodum* (kindly provided by Dr. Julian Chen at Institute of Plant Protection, Chinese Academy of Agricultural Sciences) were collected from healthy rice seedlings and starved for 2 hours before placing on the infected wild rice plants [67]. After 3 days of feeding, five aphids were carefully collected from the wild rice plants using a brush and transferred to each healthy rice seedling. Nonviruliferous aphids from healthy control rice were used as controls.

### Transmission electron microscopy

The rice samples were fixed with 2.5% glutaraldehydein at 4 °C overnight, post-fixed with 1% osmium tetroxide for 1.5 h at room temperature, dehydrated with a series of different concentrations of ethanol, and then embedded with Spurr low viscosity embedding resin at 72 °C for 12 h [68]. Ultrathin sections of rice leaves were prepared with an ultramicrotome (EM UC7), double stained with 2% uranyl acetate and 3% lead citrate, and observed under an electron microscope (FEI TECNAI G2 SPIRIT).

### Development of RITV CP antisera

The CP gene of RTIV was cloned into the prokaryotic expression vector pET30a, and the resulting recombinant plasmid was transformed into *E. coli* BL21 bacteria. The CP protein (about 22 kDa) was purified after induced expression at 15°C by IPTG. The purified protein was sent to Zoonbio Biotechnology (Nanjing, Jiangsu province, China) for antiserum production in New Zealand white rabbits.

### Construction of the full-length cDNA clone of RTIV

The 5’ and 3’ terminal sequences of RTIV were obtained by RACE and DNA sequencing using the rapid-amplification of cDNA ends kits (Invitrogen, USA). The full length genomic RNA of RTIV was converted into DNA by RT-PCR. The pCass4-RZ-RTIV infectious clone was constructed by double digestion of pCass4-RZ plasmid [51] with *Stu* I and *BamH* I, and ligation of the virus full length into the corresponding sites of pCass4-RZ. All the plasmids transformed into *Escherichia coli* JM109 and *Agrobacterium tumefaciens* EHA105 were confirmed by DNA sequencing. All primers used are listed in Table S3.

### *Agrobacterium* infiltration and inoculation

Positive *Agrobacterium* colony was cultured in Lysogeny broth fluid medium at 28°C, pelleted and resuspended in an induction buffer containing 10mM MES, 10mM MgCl_2_, 150mM acetosyringone. The *Agrobacterium* cultures were diluted to OD600 = 0.8, and then infiltrated into *N. benthamiana* leaves as described [51]. For inoculating rice plants, *Agrobacterium* strains were cultured in solid medium at 28°C until growing full culture dish and the bacteria were punctured into the crown meristematic region, leaves and stem with needle [55].

### VSR activity analysis

For the transient expression of virus gene in N. benthamiana, the RTIV gene of P0, CP, MP were amplified and introduced into pCAMBIA1300-MYC vector, which was digested with Kpn I and Sal I. All the plasmids were confirmed by DNA sequencing, transformed into Escherichia coli JM109 and A. tumefaciens EHA105. For the VSR activity test analysis, the co-infiltration A. tumefaciens (with pGD-GFP and viral genes) cultures were mixed before infiltration, and three lower leaves of N. benthamiana line 16c plants were infiltrated. GFP was visualized under ultraviolet.

### Western blot analysis

The plant tissues were collected and homogenized with protein extraction buffer (150 mM NaCl, 5 mM MgCl_2_, 20 mM Tris-HCl, pH 7.5, 5 mM dithiothreitol, 2% β-mercaptoethanol) at a ratio of 1:2 (wt/vol). Proteins were separated on sodium dodecyl sulphate (SDS) polyacrylamide gels and transferred onto Nitrocellulose membranes. The CP, Myc or AGO1a antibody was diluted at 1: 2,000, 1: 5,000 or 1: 5,000, and a peroxidase conjugated goat anti-rabbit IgG (Abmart) was used as the secondary antibody diluted at 1: 3,000.

## Acknowledgements

This work was supported by the Natural Science Foundation of China (Grant no. 31870146 to Z.G. and Grant no. 32030087 and 31871927 to Q.W.), Natural Science Foundation of Fujian Province (grant no 2020J02014 to Z.G.) and Fujian Province Hundred-Talent grant to Z.G.

## Author contribution

W-K. Y, Y. Z., W-C. and L, C-W. Z designed and performed experiments, analyzed data; S.-W. D. ZX. G. conceived, designed, and supervised the study, and wrote the paper; Q-F. W. conceived the study, supervised bioinformatics data and wrote the paper; J. B. and Z-Q. C performed experiments and analyzed data; Y-H. H and J-G. W. conceived and designed the study; R-B. L and B-S. C. provided wild rice resources and intellectual input to the study; Z-K. Z., D-L. Y. and L-H. X. provided intellectual input to the study. All authors revised and provided feedback for the final version of the paper.

## Data and materials availability

All data needed to evaluate the conclusions in the paper are present in the paper and/or the Supplementary Materials. All newly generated materials are available upon request.

## Supplementary Figures with legend

**Fig. S1.**
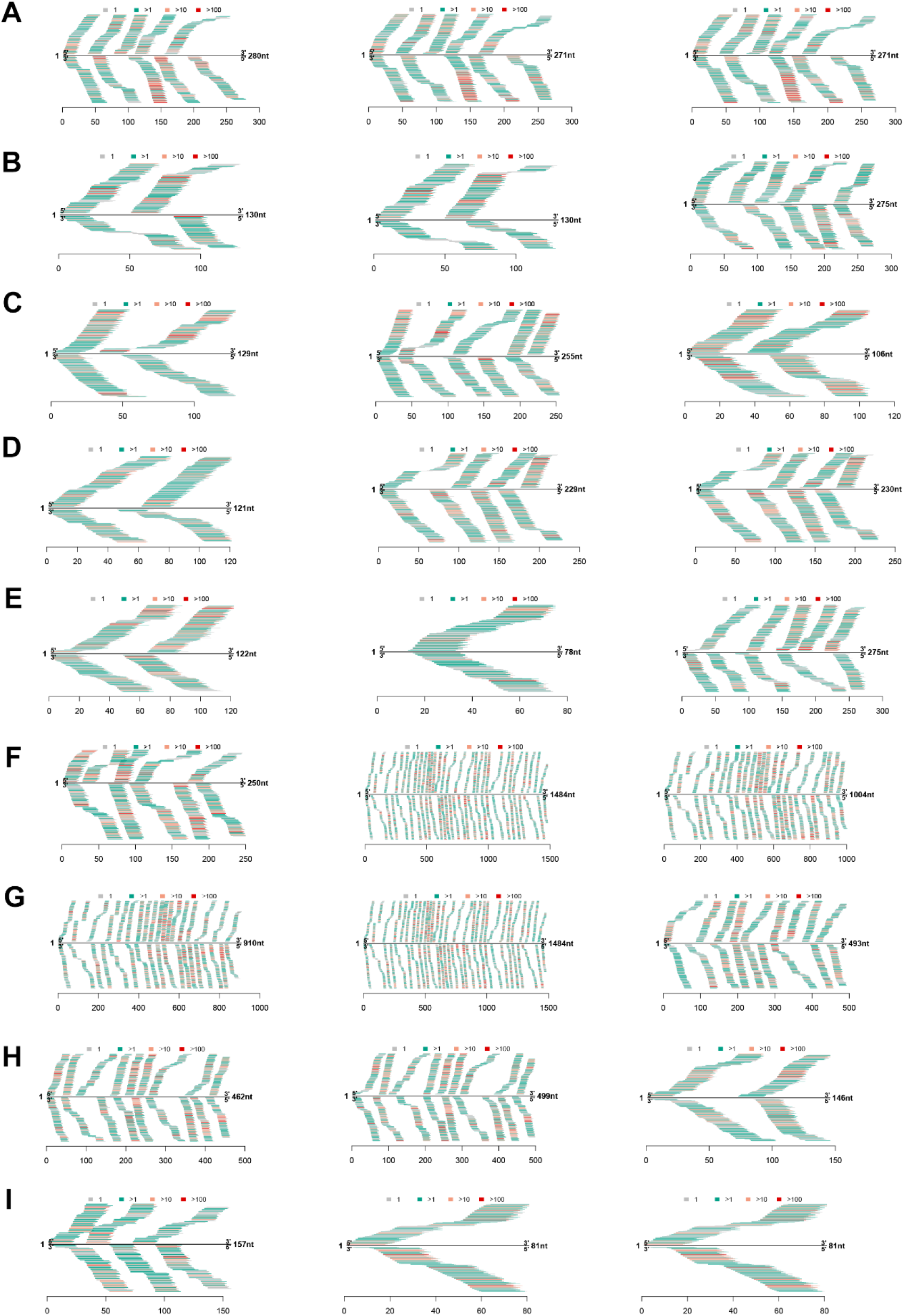
(A-I) Single RTIV contig using vdSAR based on small RNA-seq of wild rice pool.

**Fig. S2.**
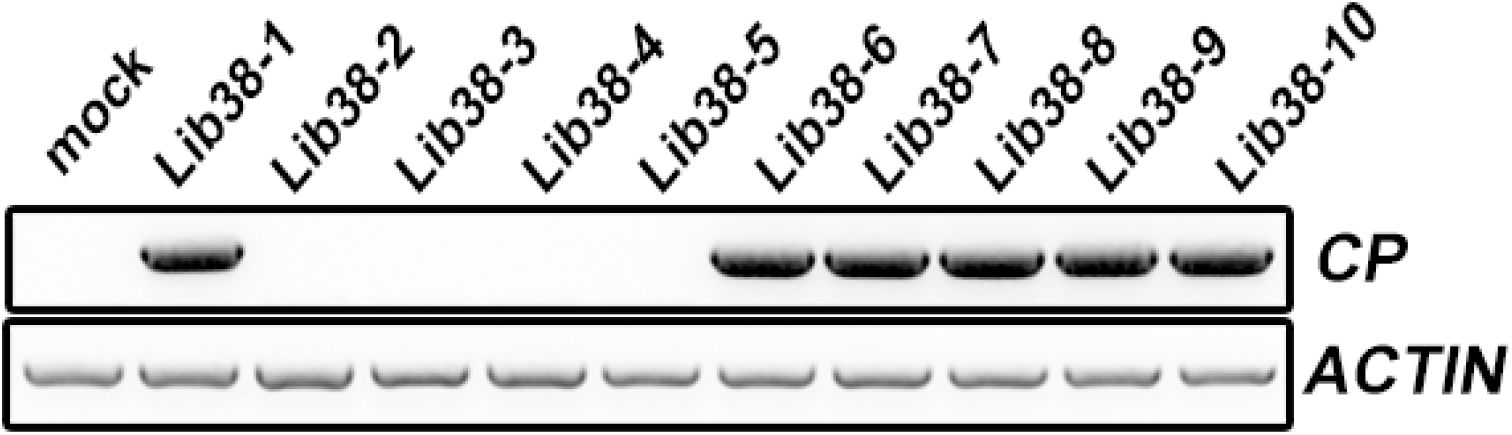
The detection of wild rice individual infected in library 38 by RT-PCR using coat protein (CP) gene-specific primers. Wild rice individual without RTIV was used as mock. Actin was used as an internal reference.

**Fig. S3.**
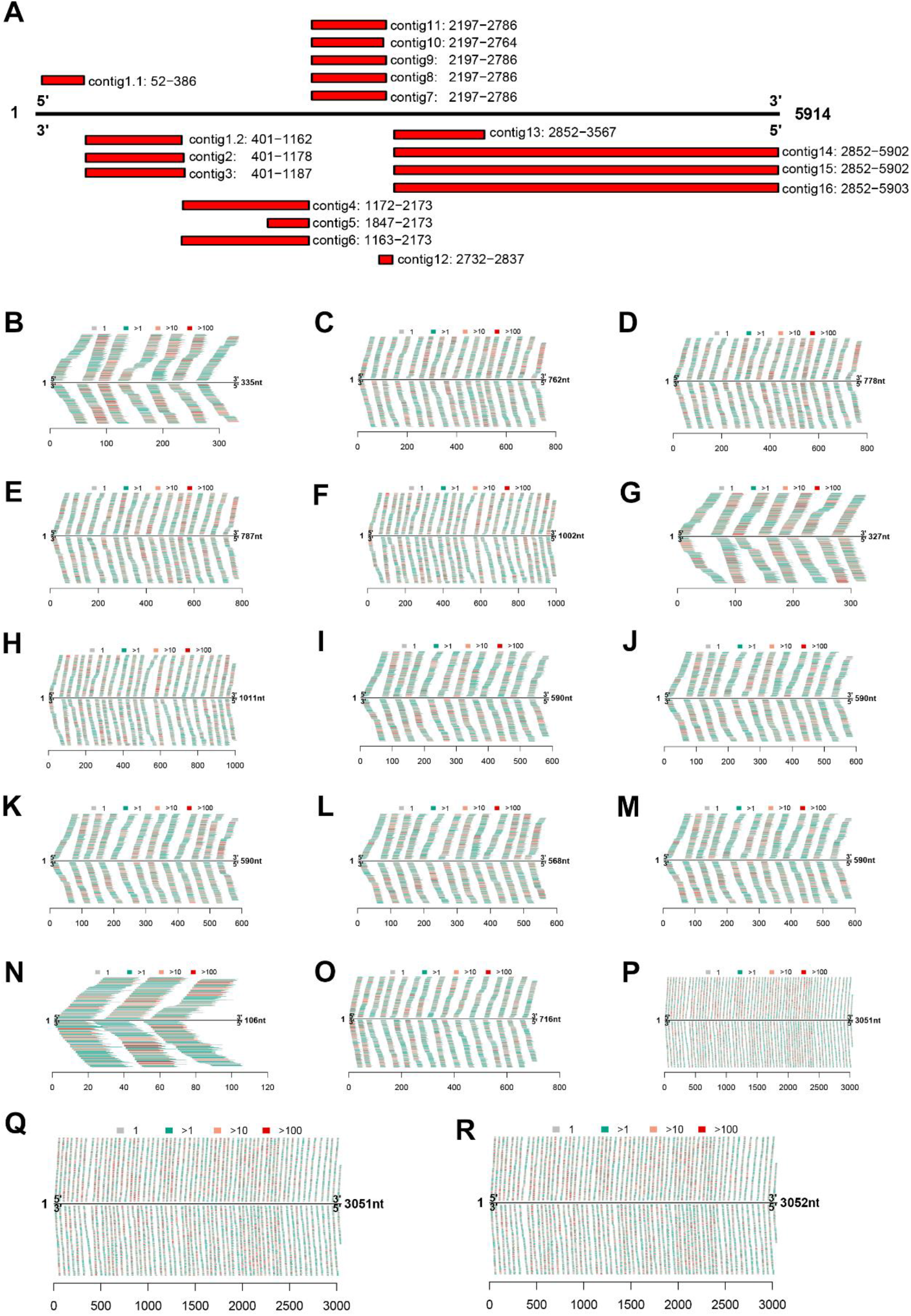
Assembled RTIV contigs mapped to RITV. **(A-R)** Each contig assembled using vdSAR based on small RNA-seq of Colony no.9 plants.

**Fig. S4.**
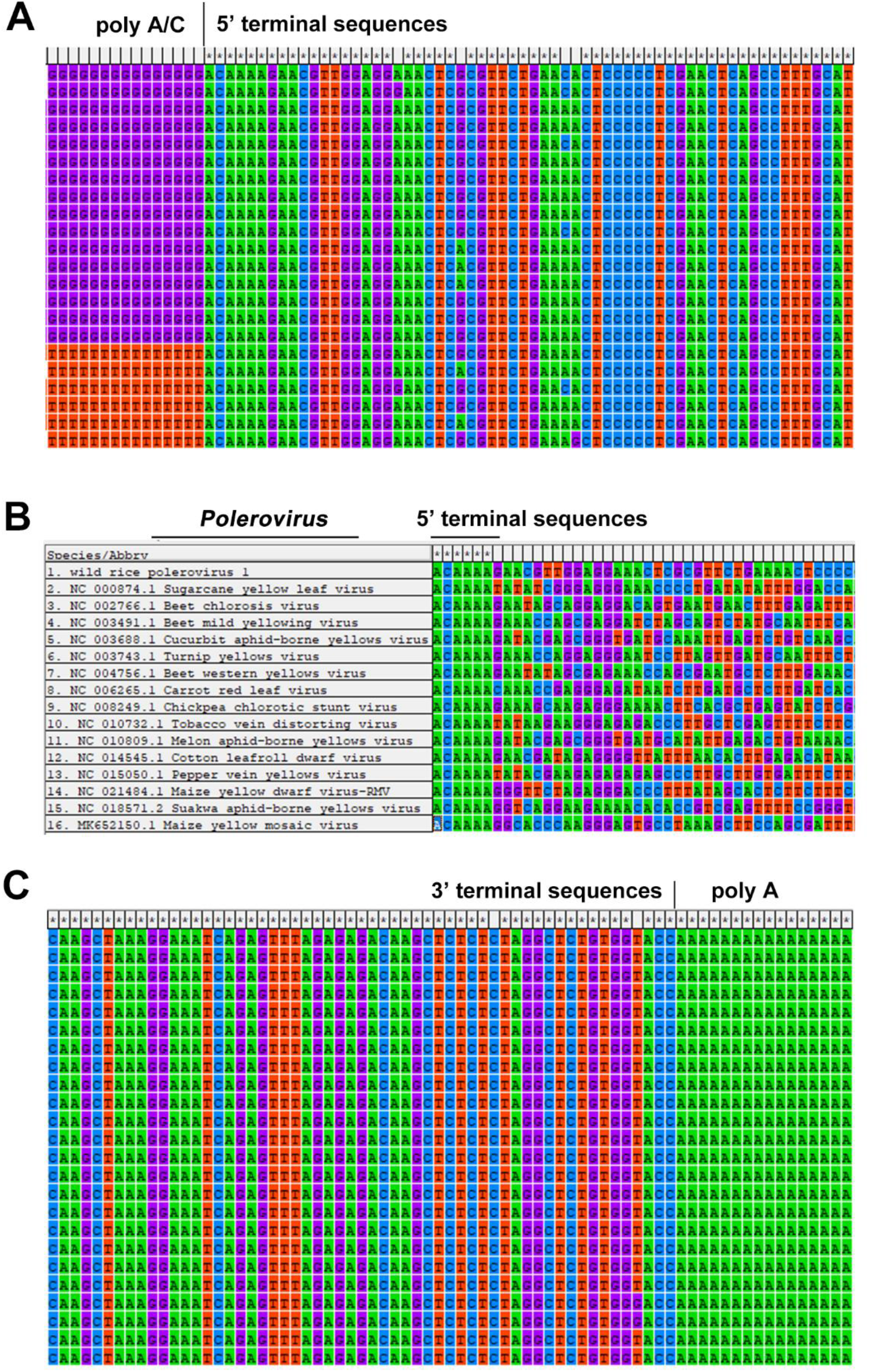
Determination of RTIV terminal sequences by 5’and 3’-RACE. **(A)** The 5’ terminal sequences were determined by 5’ RACE through adding polyA and polyC. **(B)** The 5’ terminal conserved sequences in different species of *Polerovirus*. **(C)** The 3’ terminal sequences were determined by 3’ RACE through adding polyA.

**Fig. S5.**
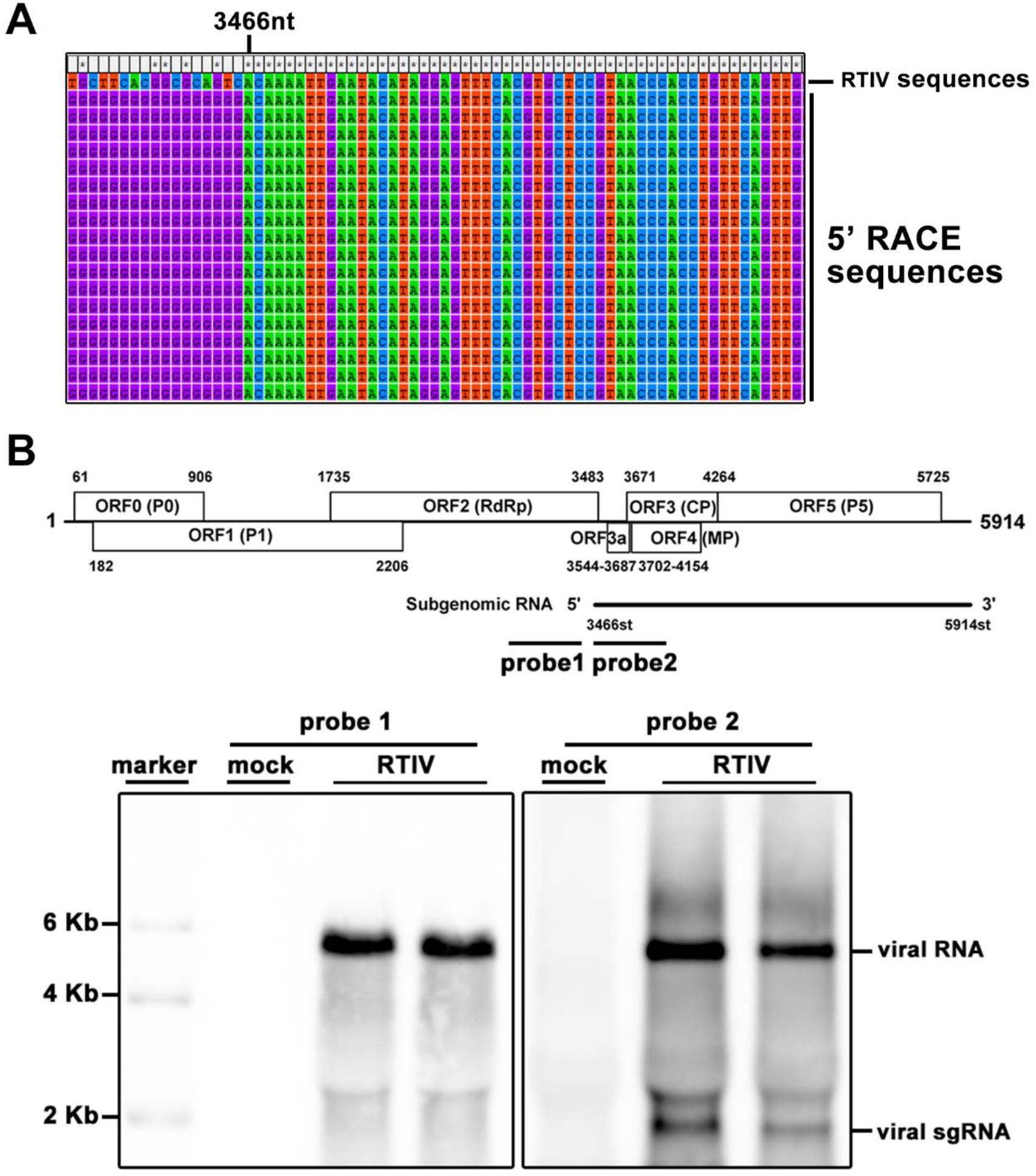
Identification of RTIV subgenomic RNA. **(A)** The 5’ terminal sequences of subgenomic RNA were determined by 5’ RACE through adding polyC. **(B)** Schematic representation of the RTIV genome and subgenomic RNA (up panel), and Northern blot confirmation of the subgenomic RNA in wild rice plants using two different probes shown in up panel (down panel).

**Fig. S6.**
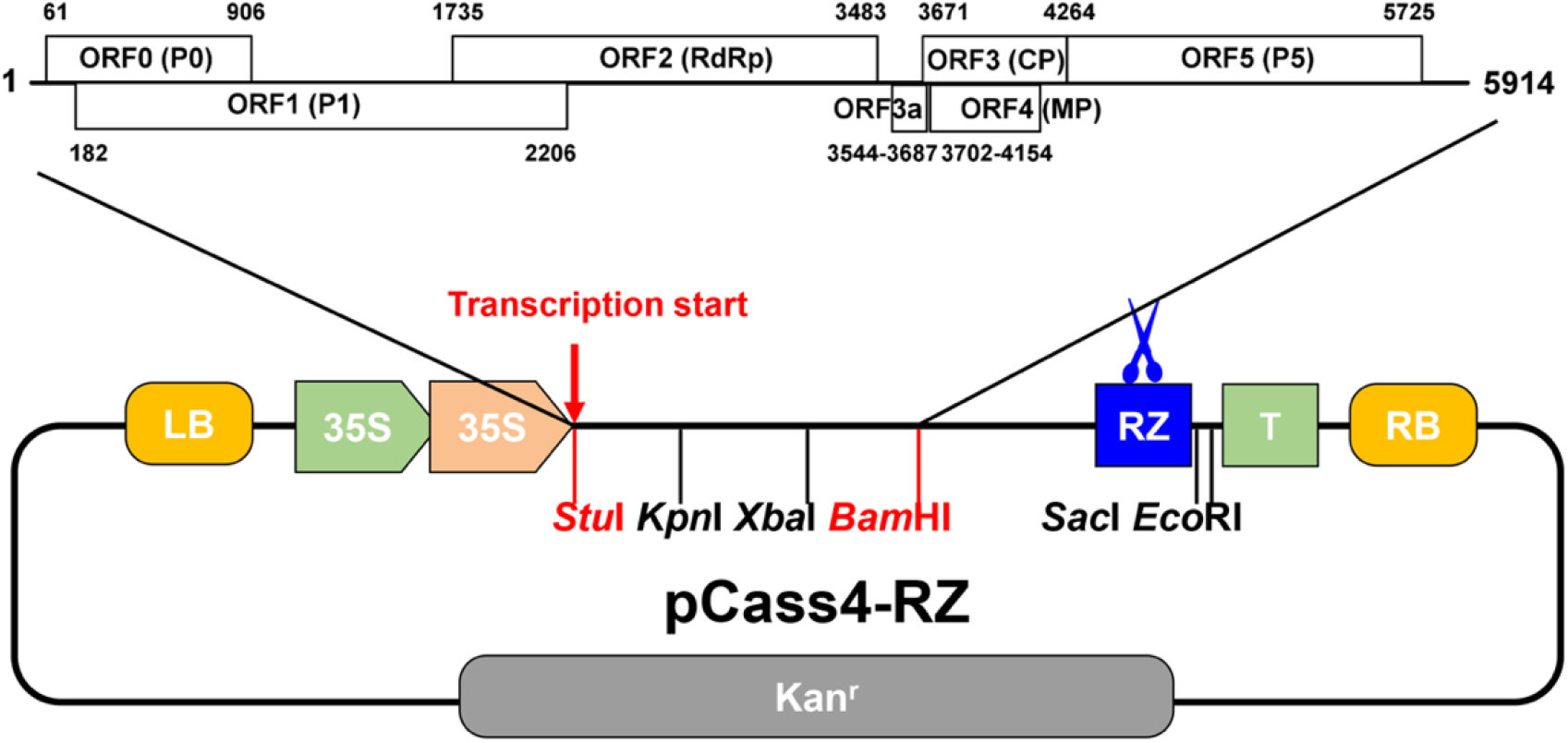
Diagram to construct pCass4-RZ-RTIV molecular clone. Full-length of RTIV was amplified and ligated to pCass4-RZ vector between StuI and BamHI sites. Primers used for vector construction listed in Table S3.

**Fig. S7.**
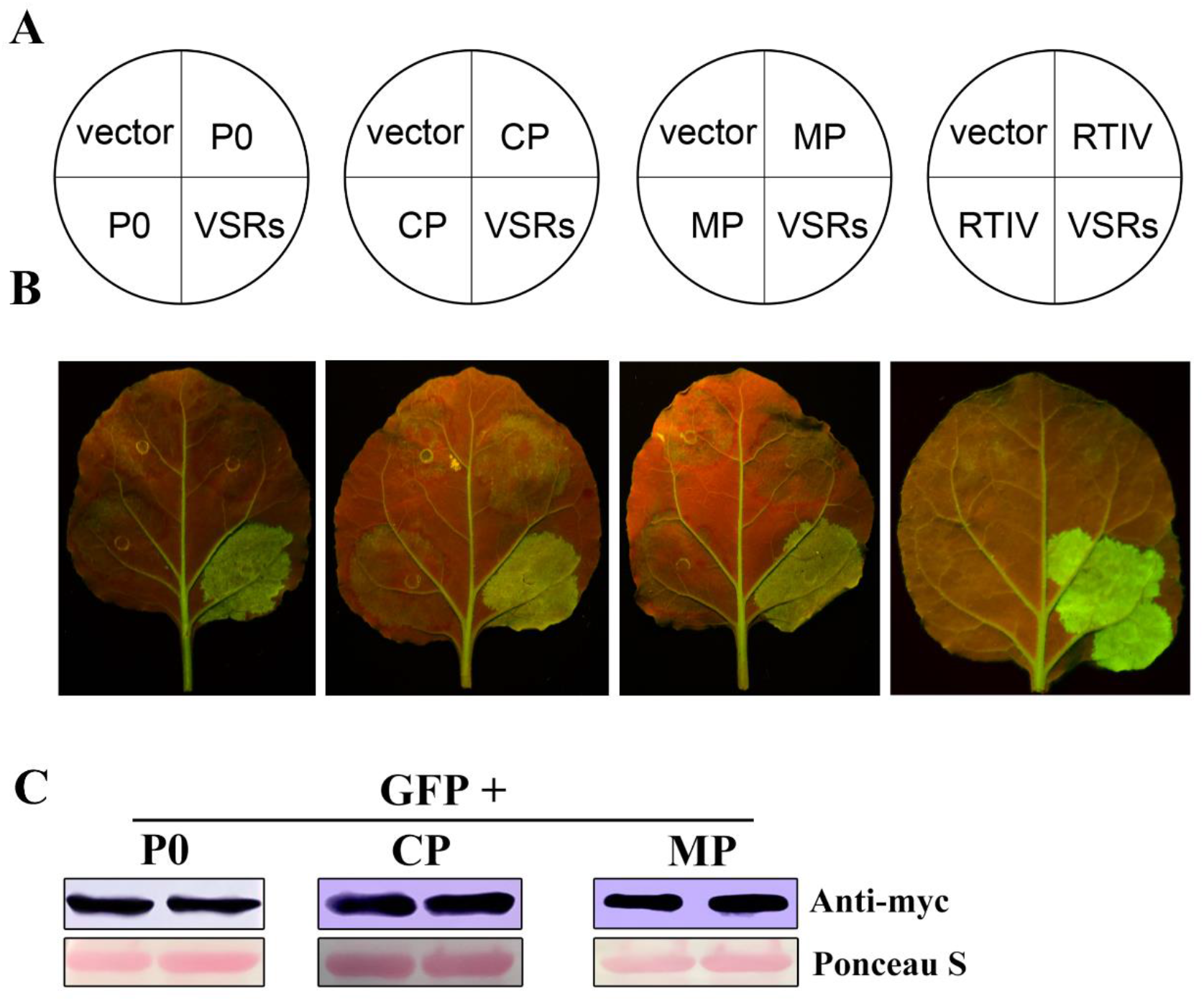
Examination of viral suppressors of RNA-silencing (VSR) in RTIV. **(A)** Schematic representation of the Agrobacterium infiltration. **(B)** 16C transgenic N. benthamiana were respectively co-infiltrated with GFP and P0, CP, MP or full length of RTIV. Empty vector and VSRs were used as negative and positive control, respectively. GFP was visualized under ultraviolet 5pdi. (C) The expression of P0, CP or MP tagged with Myc was confirmed by Western blot, Ponceau S used as loading control.

**Fig. S8.**
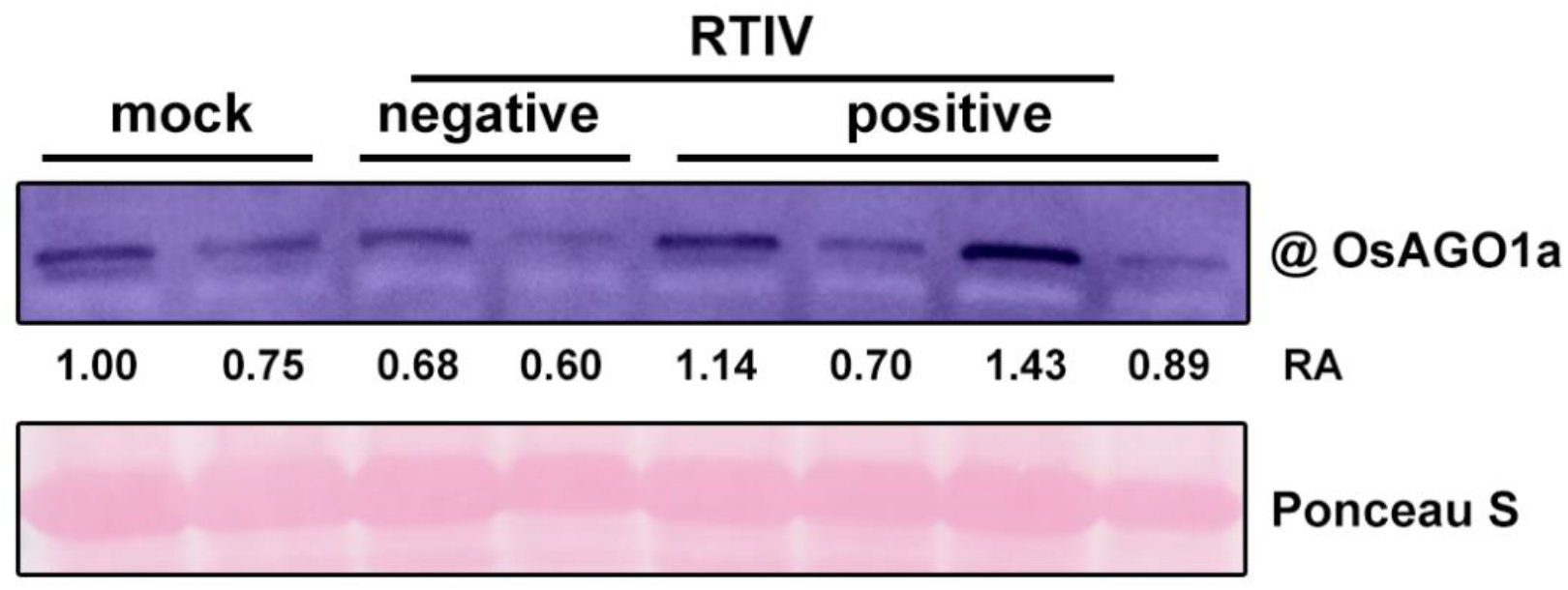
RTIV infection does not affect expression level of AGO1a protein in rice plants. Western blot detection of AGO1a protein accumulation in RTIV-positive and RTIV-negative Zhonghua-11 rice plants at ~30dpi compared to mock Zhonghua-11 rice plants, RA is the relative accumulation of AGO1a normalized to loading control, the same membrane was stained for Ponceau S to show loading control.

**Fig. S9.**
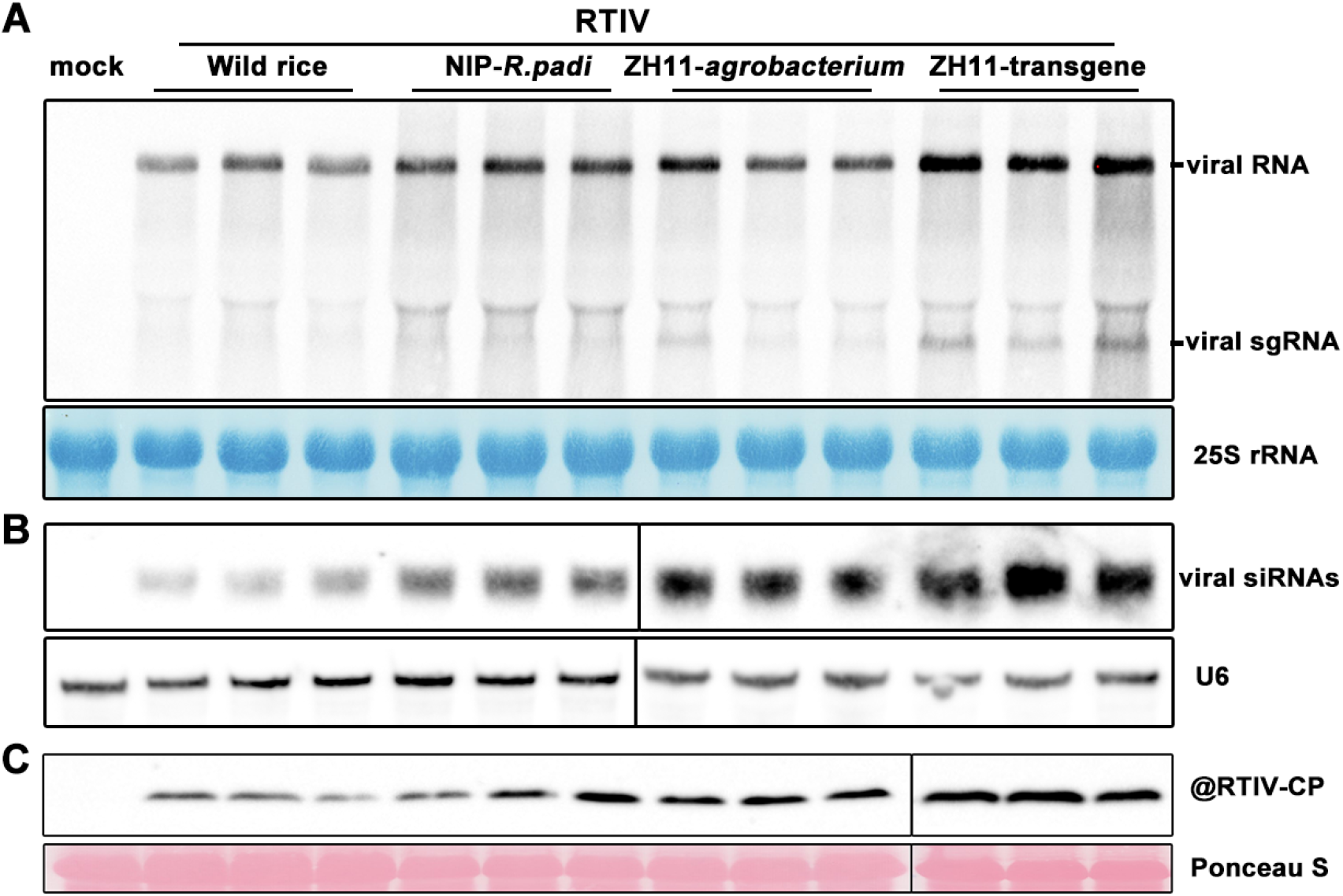
Comparison of viral RNA or vsiRNA accumulation in Wild rice plants, NIP infected RTIV through aphid, ZH11 mechanically infected RTIV molecular clone or RTIV-amplicon transgenic ZH11. **(A)** Northern blot detection of viral genomic RNAs accumulation, 25S rRNA in the same membrane was stained to show equal RNAs loading. **(B)** Northern blot detection of viral siRNAs accumulation, the same membrane was probed for U6 RNA to show equal small RNAs loading. **(C)** Western blot detection of viral coat protein accumulation, the same membrane was stained for ponceau to show equal protein loading.

**Fig. S10.**
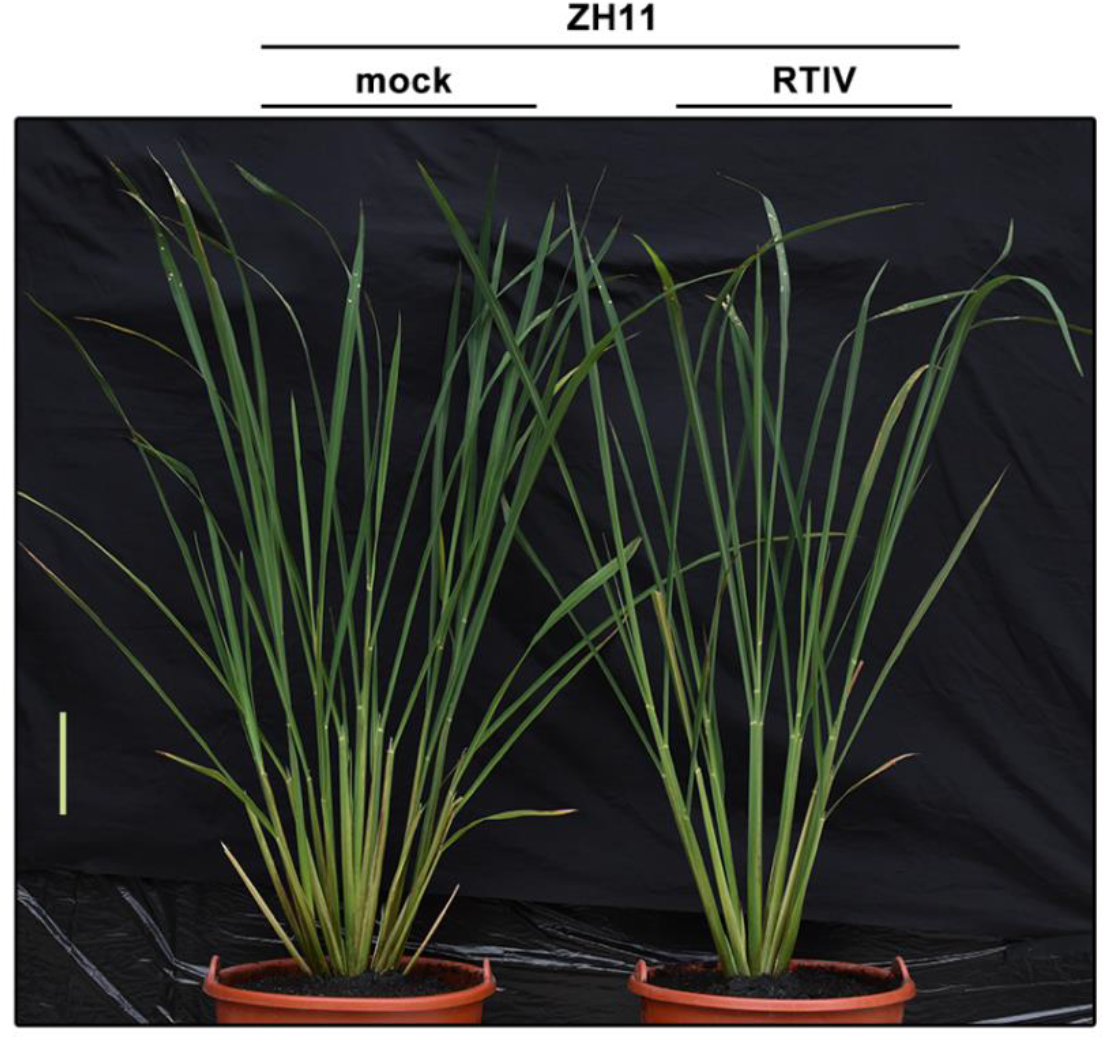
Low-tillering disease symptom in rice cultivar ZH11 after the infection of RTIV transmitted by aphid. RTIV was acquired from RTIV-positive transgenic rice plants by aphid *R. Padi* and transmitted to healthy wildtype ZH11. Plants were photographed at ~45 dpi. Scale bar: represent 10 cm.

